# Overlapping and distinct cis-acting requirements for *oskar* mRNA localization pathways

**DOI:** 10.1101/2022.04.04.487058

**Authors:** Catherine E. Eichler, Michelle E. Grunberg, Elizabeth R. Gavis

## Abstract

Localization of *oskar* mRNA to the posterior of the *Drosophila* oocyte is essential for abdominal patterning and germline development. *oskar* localization is a multi-step process involving temporally and mechanistically distinct transport modes. Numerous cis-acting elements and trans-acting factors have been identified that mediate earlier motor-dependent transport steps leading to accumulation of *oskar* at the posterior. Little is known, however, about what features of *oskar* are required for the later localization phase, that occurs by diffusion with local entrapment and results in the accumulation of large *oskar* ribonucleoprotein granules by the end of oogenesis. Here we test whether cis- and trans-acting requirements for kinesin-dependent *oskar* transport, the structured *oskar* spliced localization element (SOLE) and the exon junction complex (EJC), are also required for late-phase localization. In contrast to kinesin-dependent motility, late-phase localization is independent of the EJC and depends not only on the structure but also the sequence of the SOLE. Late-phase localization also requires the *oskar* 3’ UTR and we show that the 3’ UTR is sufficient for ribonucleoprotein granule selectivity.

## INTRODUCTION

Localization of mRNAs to subcellular domains plays an important role in generating morphological and functional asymmetry through local protein production. Localization of *oskar* (*osk*) mRNA to the posterior of the *Drosophila* oocyte and its onsite translation is essential for formation of the germ plasm, the specialized embryonic cytoplasm required for abdominal patterning and germ cell formation during embryogenesis (Ephrussi et al., 1991; Kim-Ha et al., 1991; Lehmann and Nusslein-Volhard, 1986; Rongo et al., 1995). The trafficking of RNAs to different subcellular locations is controlled by cis-acting RNA sequences and/or structures that most commonly reside within 3’ untranslated regions, and proteins that recognize these localization elements. These proteins may interface with cellular transport machinery or they may facilitate association with cellular structures or organelles, including phase transitioned condensates (Das et al., 2021; Martin and Ephrussi, 2009). Multiple cis-acting elements along with a coterie of trans-acting factors, contribute to the multi-step process of *osk* localization throughout oogenesis.

Drosophila oogenesis proceeds through 14 morphologically defined stages; during the first 10 stages the oocyte is accompanied by 15 sister nurse cells that support its growth and development by providing maternal mRNAs such as *osk*, proteins, and organelles. *osk* is transported from the nurse cells to the oocyte by dynein; this transport requires multiple sequence elements and structures in the 3’ UTR (Jambor et al., 2014; Ryu et al., 2017). One of these, called the oocyte entry signal (OES) links *osk* RNA to dynein by binding to the RNA-binding protein Egalitarian and the adaptor Bicaudal-D (Dienstbier et al., 2009; Goldman et al., 2019; Jambor et al., 2014). Throughout stage 8 to 10, *osk* travels to the posterior of the oocyte by kinesin-mediated transport (Brendza et al., 2000; Zimyanin et al., 2008). This transport requires an unusual localization element, a stem-loop structure called the Spliced *oskar* Localization Element (SOLE) (Ghosh et al., 2012). The SOLE comprises the last 18 nucleotides of the first *osk* exon and the first 10 nucleotides of the second *osk* exon and therefore splicing of the first *osk* intron is crucial for localization. Mutational analysis showed that SOLE function depends on its structural integrity rather than on its sequence. Splicing of the first *osk* intron also plays a role in *osk* localization through recruitment of the exon junction complex (EJC) (Ghosh et al., 2012). Both the SOLE and EJC proteins including eIF4AIII, Y14, Mago Nashi, and Barentsz are required for *osk* transport (Hachet and Ephrussi, 2001; Mohr et al., 2001; Newmark and Boswell, 1994; Palacios et al., 2004; van Eeden et al., 2001) and live imaging studies showed that the EJC/SOLE activates kinesin motility leading to accumulation of *osk* at the posterior (Gaspar et al., 2017; Ghosh et al., 2012).

In addition to its role in nurse cell-to-oocyte transport, the *osk* 3’ UTR contributes in multiple ways to posterior transport within the oocyte (Kim-Ha et al., 1993). In the oocyte, *osk* multimerizes, forming ribonucleoprotein particles (RNPs) containing up to 4 *osk* mRNAs (Little et al., 2015). Two proteins, Bruno and polypyrimidine tract-binding protein/hnRNP I (PTB), bind to multiple sites in the *osk* 3’ UTR and promote *osk* oligomerization to form higher order complexes, although these are associated with repression of *osk* translation, rather than localization (Besse et al., 2009; Chekulaeva et al., 2006). In addition, a short sequence at the tip of the OES called the dimerization domain promotes *osk* dimerization, allowing transgenic reporter RNAs with the dimerization domain, but lacking the SOLE, to “hitchhike” to the posterior on endogenous wild-type *osk* RNA (Hachet and Ephrussi, 2004; Jambor et al., 2011; Kim-Ha et al., 1993). Whether hitchhiking contributes to localization of endogenous *osk* remains to be determined, however.

Finally, the *osk* 3’ UTR contains binding sites for the double-stranded RNA-binding protein Staufen (Stau) (Mohr et al., 2021). Stau associates with *osk* RNPs upon their entry to the oocyte and remains colocalized with *osk* throughout the rest of oogenesis (Ephrussi et al., 1991; Kim-Ha et al., 1991; Little et al., 2015; Micklem et al., 2000; St Johnston et al., 1991). Stau acts in part by counteracting cortical anchoring of *osk* that occurs upon its entry into the oocyte, permitting its transport to the posterior by kinesin (Mohr et al., 2021; St Johnston et al., 1991; Zimyanin et al., 2008). Stau is also required for optimal kinesin activity and for the selective activation of *osk* translation at the posterior and *osk* anchoring (Micklem et al., 2000).

Accumulating at the posterior, Osk protein establishes a posterior domain of germ plasm assembly. Osk recruits additional proteins including the helicase Vasa and initiates formation of phase-transitioned condensates called germ granules, which will ultimately be inherited by the germ cell progenitors during embryogenesis (Lehmann, 2016). At the end of stage 10 of oogenesis, the nurse cells initiate a cell death program and extrude their contents en masse into the oocyte. Just prior to this nurse cell dumping, the microtubules reorganize into parallel bundles at the cortex of the oocyte allowing mixing of the oocyte and nurse cell contents in a process called ooplasmic streaming (Becalska and Gavis, 2009; Spradling, 1993). A subset of mRNAs circulating within the oocyte during this time become entrapped by the prelocalized germ granule protein assemblies (Forrest and Gavis, 2003; Little et al., 2015). Among these is *nanos* (*nos*) mRNA, whose translation requires Osk and is crucial for both embryonic abdominal patterning and germ cell development (Asaoka et al., 1998; Gavis and Lehmann, 1994; Lehmann and Nusslein-Volhard, 1991; Wang and Lehmann, 1991). For many germ granule localized mRNAs including *nos*, elements contained within 3’ UTRs are sufficient to direct their entrapment (Eagle et al., 2018; Gavis et al., 1996; Little et al., 2015; Rangan et al., 2009). Notably, additional *osk* mRNA accumulates at the posterior during this later period through a similar diffusion and entrapment mechanism rather than by microtubule transport (Glotzer et al., 1997; Sinsimer et al., 2011). *osk* RNPs remain distinct from germ granules and grow in number and size, accumulating up to 250 *osk* mRNAs (Little et al., 2015). We refer to these large *osk* assemblies as founder granules, to distinguish them from germ granules. This late phase of *osk* accumulation amplifies the germ plasm, allowing production of additional Osk protein, enlargement of germ granules, and accumulation of germ granule mRNAs (Niepielko et al., 2018; Sinsimer et al., 2011; Snee et al., 2007). Since the quantity of germ plasm formed, and consequently the quantities of the abdominal determinant Nos and germ cells produced in the embryo, depends directly on the amount of *osk* mRNA localized during oogenesis (Ephrussi and Lehmann, 1992; Gavis and Lehmann, 1994; Smith et al., 1992), this amplification is crucial for abdominal patterning and reproduction. Notably, failure to accumulate *osk* mRNA and protein at late-stages of oogenesis results in loss of abdominal segments and germ cell progenitors (Sinsimer et al., 2011; Snee et al., 2007).

Which features of *osk* mediate this late localization phase and whether they are distinct from the signals that mediated the earlier microtubule-dependent steps remains unknown. Here we begin to answer this question by analyzing the behavior of *osk* transgenes with mutations or deletions that affect the *osk* SOLE and 3’ UTR. We show that, in contrast to localization during stages 8-10, both the structure and the sequence of the SOLE, but not the EJC, is necessary for late-phase localization. Late-phase localization also requires the *osk* 3’ UTR and by swapping the *nos* and *osk* 3’ UTRs, we show that each is sufficient to target RNA to the appropriate granule.

## RESULTS

### Late phase *osk* mRNA localization does not require the EJC

We first tested whether elements involved in the active transport of *osk* to the posterior during stages 8-10 also function in *osk* localization during late oogenesis. We generated flies expressing a genomic *osk* transgene tagged with a superfolder *gfp* (*sfgfp*) sequence (*osk-sfgfp*), as well as a version with a deletion of the first two introns to disrupt splicing-dependent deposition of the EJC adjacent to the SOLE (*oskΔi1,2-sfgfp*) (Figure 1A). The *sfgfp* sequence tag allows for detection of the transgenic mRNA by fluorescence in situ hybridization (FISH) in the context of endogenous *osk*. Since the *oskΔi1,2-sfgfp* transgenic mRNA is not expected to localize on its own by kinesin-dependent transport, we reasoned that the presence of endogenous *osk*, and consequent production of Osk protein, would be necessary for the initial establishment of germ plasm and for the retention of *osk* mRNA during late stages of oogenesis. The *osk-sfgfp* and *oskΔi1,2-sfgfp* were expressed at comparable levels as determined by RT-qPCR (Figure S1).

**Figure 1.**
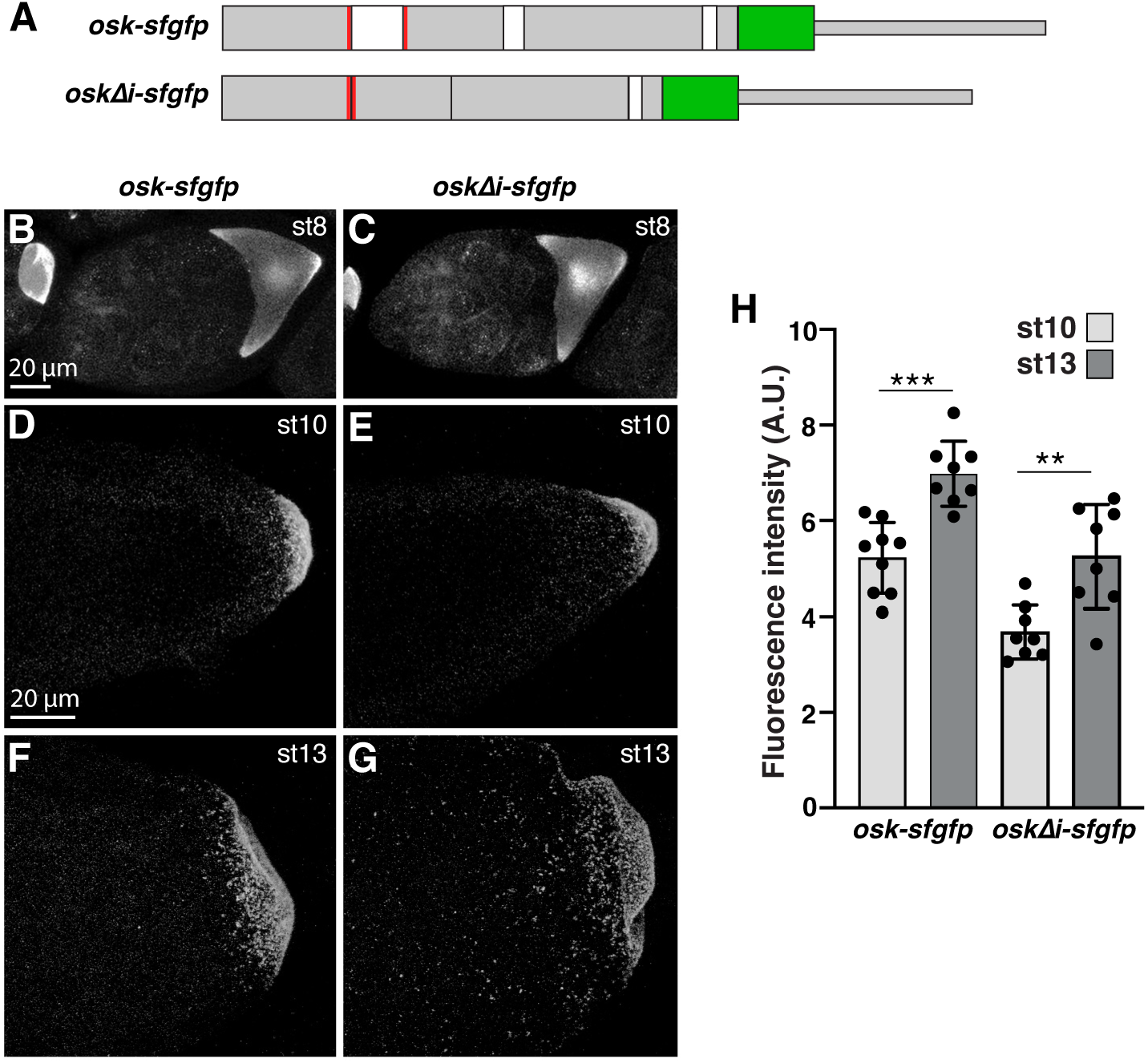
The EJC is not required for late-phase *osk* localization. (A) Structure of *osk-sfgfp* and *oskΔi-sfgfp* transgenes. Grey boxes: *osk* exonic sequences; white boxes: *osk* introns; green boxes: *sfgfp*; red lines: sequences that create the SOLE; thinner boxes indicate 3’ UTRs. (B-G) Confocal z series projections of transgenic stage 8 egg chambers (B, C), stage 10 oocytes (D, E), and stage 13 oocytes. The entirety of the germ plasm was captured. Transgenic RNAs were detected by FISH using probes for *sfgfp*. Probes were labeled with 647 fluorophores to avoid detecting fluorescence from Osk-sfgfp protein. (J) Quantification of total localized fluorescence signal intensity. n = 8-9 oocytes each. Values are mean ± standard deviation. ** p < 0.01, *** p < 0.001 as determined by Students t-test.

Quantification of the total localized *sfgfp* fluorescence intensity at stage 10 showed that in the presence of wild-type *osk*, both the *osk-sfgfp* and *oskΔi1,2-sfgfp* mRNAs could localize during mid-oogenesis (Figure 1B-E, H). However, localization of the *oskΔi1,2-sfgfp* mRNA was less efficient than the *osk-sfgfp* mRNA. This is consistent with the finding that without EJC deposition on *osk* mRNA, transport efficiency of *osk* RNPs is reduced (Ghosh et al., 2012) and suggests that the *oskΔi1,2-sfgfp* mRNA only localizes by hitchhiking with endogenous *osk* using the dimerization domain in the 3’ UTR (Jambor et al., 2011). By stage 13, the localized fluorescence intensities of the *osk-sfgfp* and *oskΔi1,2-sfgfp* mRNAs increased by 34% and 43%, respectively (Figure F-H). Therefore, the EJC is not required for late-phase *osk* localization.

### Late phase *osk* localization requires the SOLE UA-rich proximal stem sequence

Although the *oskΔi1,2-sfgfp* mRNA is not bound by the EJC at the first exon-exon junction, it can form the SOLE. To test whether the SOLE influences late phase *osk* localization independently of the EJC, we generated a version of the *osk-sfgfp* transgene with the SOLE proximal stem sequence substituted by *lacZ* sequence (*SOLE*^*PS-Lz*^; Figure 2A). This mutation was previously shown to disrupt SOLE localization activity during mid-oogenesis without affecting EJC deposition (Ghosh et al., 2012). As expected, the *oskSOLE*^*PS-Lz*^*-sfgfp* mRNA localized to the posterior of the oocyte by stage 10 in the presence of endogenous *osk*, although with reduced efficiency compared to the *osk-sfgfp* control mRNA (Figure 2D, J). This is consistent with the reduced localization efficiency at stage 10 of the *oskΔi1,2-sfgfp* mRNA. However, in contrast to the *oskΔi1,2-sfgfp* mRNA, the *oskSOLE*^*PS-Lz*^*-sfgfp* mRNA was not further enriched by stage 13 (Figure 3E, J). Therefore, the SOLE is required for late-phase *osk* localization, and this role is independent of the EJC.

**Figure 2.**
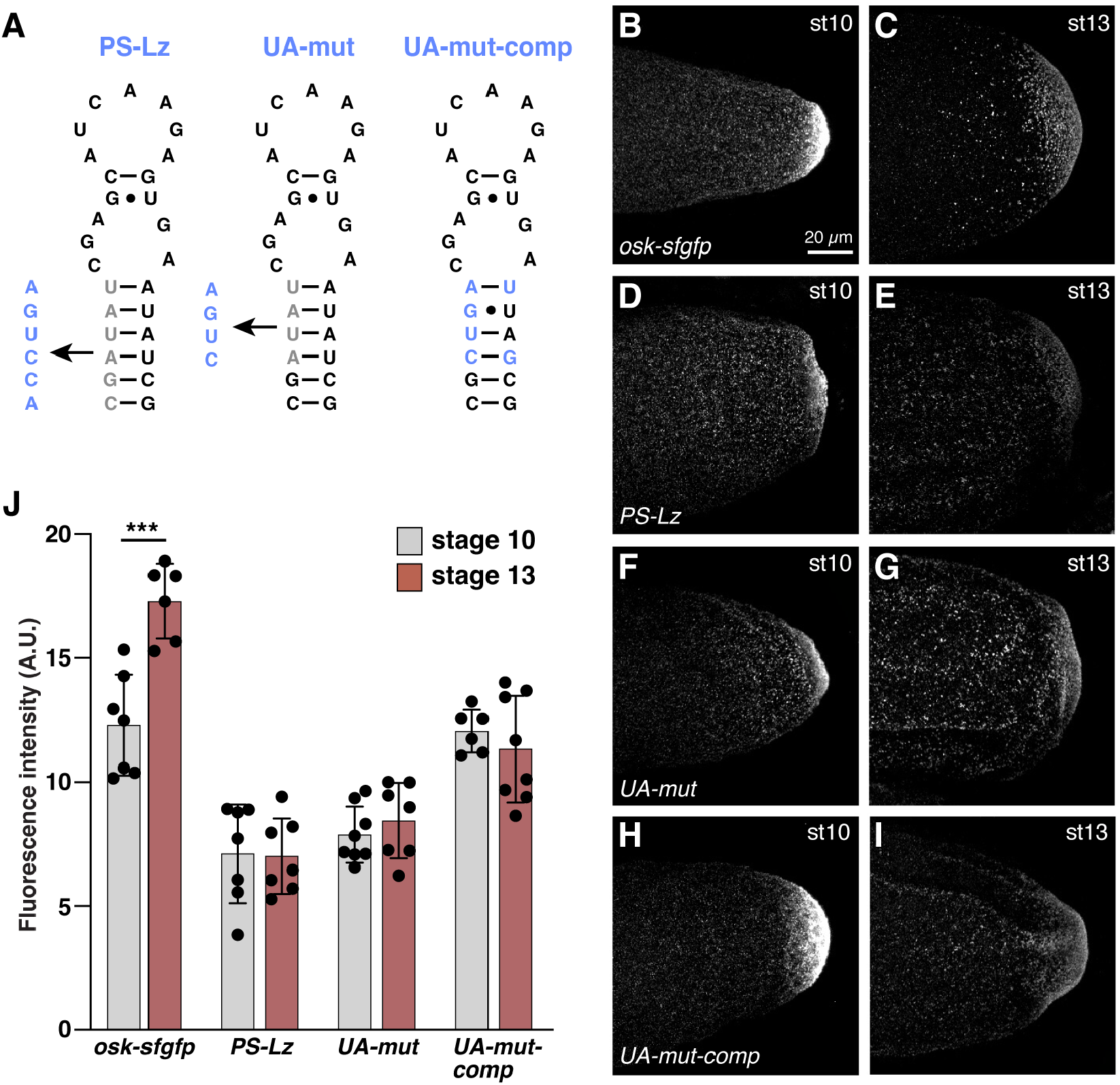
The sequence and structure of the SOLE is required for late-phase *osk* localization. (A) The secondary structure of the SOLE with nucleotides of the proximal stem that were changed indicated in gray and the new sequences shown in blue. (B-I) Confocal z series projections of transgenic stage 10 (B, D, F, H) and stage 13 (C, E, G, I) oocytes. The entirety of the germ plasm was captured. Transgenic RNAs were detected by FISH using 647-labeled probes for *sfgfp*. (J) mean ± standard deviation. *** p < 0.001 as determined by Student’s t-test.

**Figure 3.**
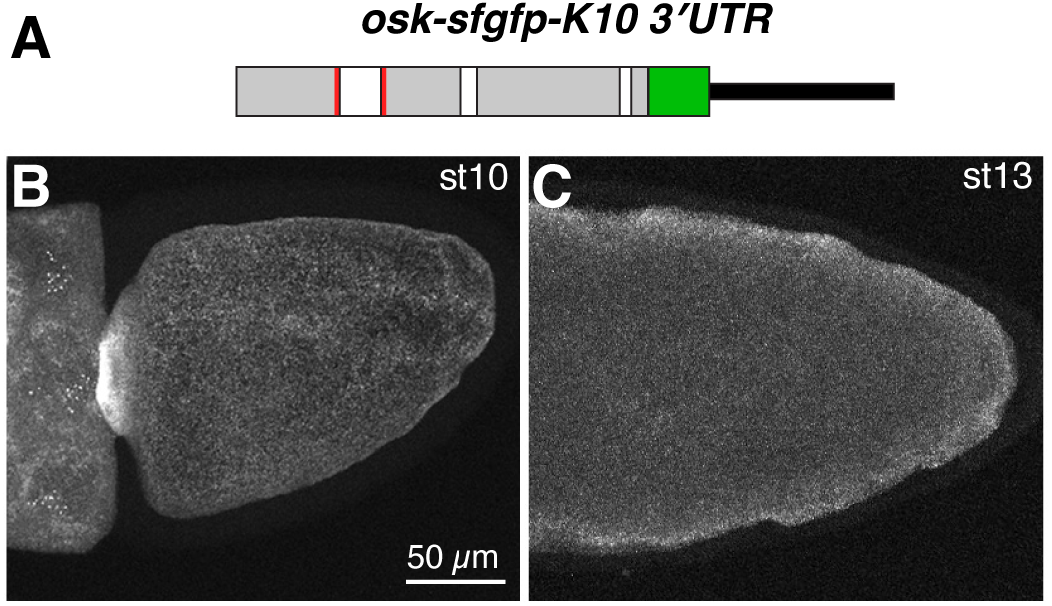
The *osk* 3’ UTR is required for late-phase localization. (A) Structure of the *osk-sfgfp-K10 3’UTR* transgene. Grey boxes: *osk* exonic sequences; white boxes: *osk* introns; green boxes: *sfgfp*; red lines: sequences that create the SOLE; thin black bar: *K10* 3’UTR. (B) Confocal z projections of transgenic stage 10 (B) and stage 13 (C) oocytes, with transgenic RNA detected by FISH. ≥10 oocytes were imaged for each stage, with similar results.

The earlier phase of *osk* localization relies on the structure, but not the sequence of the SOLE proximal stem (Ghosh et al., 2012). To determine whether this is also the case for the late phase of *osk* localization, we generated a new mutation that disrupts base-pairing of the proximal stem and is thus predicted to disrupt the regular helical structure of the SOLE (*SOLE*^*UA-mut*^) (Simon et al., 2015), as well as a compensatory mutation (*SOLE*^*UA-mut*-comp^) that restores the proximal stem base-pairing, but not the sequence (Figure 2A). Like the *oskSOLE*^*PS-Lz*^*-sfgfp* mRNA, *oskSOLE*^*UA-mut*^*-sfgfp* mRNA showed reduced localization efficiency at stage 10 in the context of endogenous *osk* and failed to enrich further at stage 13 (Figure 2F, G, J). The compensatory *SOLE*^*UA-mut*-comp^ mutation restored localization efficiency at stage 10, but did not restore late phase localization (Figure 2 H-J). This is in contrast to the earlier phase of *osk* localization (Ghosh et al., 2012). In all cases, RT-qPCR analysis confirmed that loss of enrichment was not due to low expression of the transgenic RNA (Figure S1). Thus, these data show that the UA-rich sequence of the proximal stem, but not the structure alone is required for late-phase *osk* localization.

### Late phase *osk* localization requires the *osk* 3’ UTR

In addition to the SOLE, the *osk* 3’ UTR is required for *osk* localization during stages 8-10 (Kim-Ha et al., 1993). Furthermore, the *osk* 3’ UTR of an RNA lacking the SOLE can facilitate localization by dimerizing with the 3’ UTR of wild-type endogenous *osk*, allowing the SOLE-less RNA to hitchhike to the posterior. The finding that none of the SOLE mutants tested above supported late-phase accumulation suggests that hitchhiking is not occurring at these later stages or does not result in retention of *osk* at the posterior. To test whether the 3’ UTR plays any role in late-phase localization, we generated an *osk-sfgfp* transgene (including the SOLE) in which the *osk* 3’ UTR was replaced by the *fs(1)K10* 3’ UTR (Figure 3A). *osk-sfgfp-K10_3*’*UTR* RNA was readily detectable by FISH when expressed under GAL4/UAS control. As expected, *osk-sfgfp-K10_3*’*UTR* RNA did not accumulate at the posterior of the oocyte prior to stage 10 even in the presence of wild-type *osk* (Figure 3B). Furthermore, the *osk-sfgfp-K10_3*’*UTR* did not localize by stage 13 either (Figure 3C), indicating that the *osk* 3’ UTR is required for both earlier and later phases of *osk* localization.

### The *osk* 3’ UTR is sufficient to localize RNA to founder granules

Localized *osk* mRNA forms large RNP bodies in late-stage oocytes and embryos that contain numerous *osk* transcripts and Stau protein. These founder granules are distinct from germ granules, which harbor mRNAs that will be inherited by the germ cell progenitors during embryogenesis (Eichler et al., 2020; Little et al., 2015). Studies of several germ granule mRNAs, including *nanos* (*nos*), *polar granule component*, and *germ cell-less*, have shown that their 3’ UTRs are sufficient for their accumulation in germ granules. To determine if the *osk* 3’ UTR is sufficient to direct RNA to founder granules, we generated tagged, genomic *osk* and *nos* transgenes with their 3’UTRs exchanged (Figure 4A) and asked whether the hybrid RNAs were targeted to founder granules or germ granules. The *sfgfp* and *egfp* sequences allowed the respective transgenic RNAs to be distinguished from endogenous *osk* and *nos* by FISH. *cyclin B* (*cycB*), an abundant germ granule transcript, was used as a germ granule marker and Stau as a founder granule marker. We performed dual FISH and immunofluorescence to detect the transgenic mRNAs with *sfgfp* or *egfp* probes coupled to 670 or 647 fluorophore, respectively; germ granules with *cycB* probes coupled to 565 fluorophore; and founder granules with anti-Stau and a secondary antibody coupled to 488 fluorophore. The experiments were performed using early embryos, due to the impenetrability of late-stage oocytes to antibodies. We defined germ granule-association as 670/647 fluorescence signal (transgenic mRNA) colocalized with 565 fluorescence signal (*cycB*). Because the transgenes produce GFP-labeled proteins, to eliminate any contribution from residual GFP fluorescence that overlaps with germ granules and founder granules, we defined founder granule association as 670/647 fluorescence signal colocalized with 488 fluorescence signal (Stau) that was not associated with 565 fluorescence (*cycB)*, which should represent Stau only.

**Figure 4.**
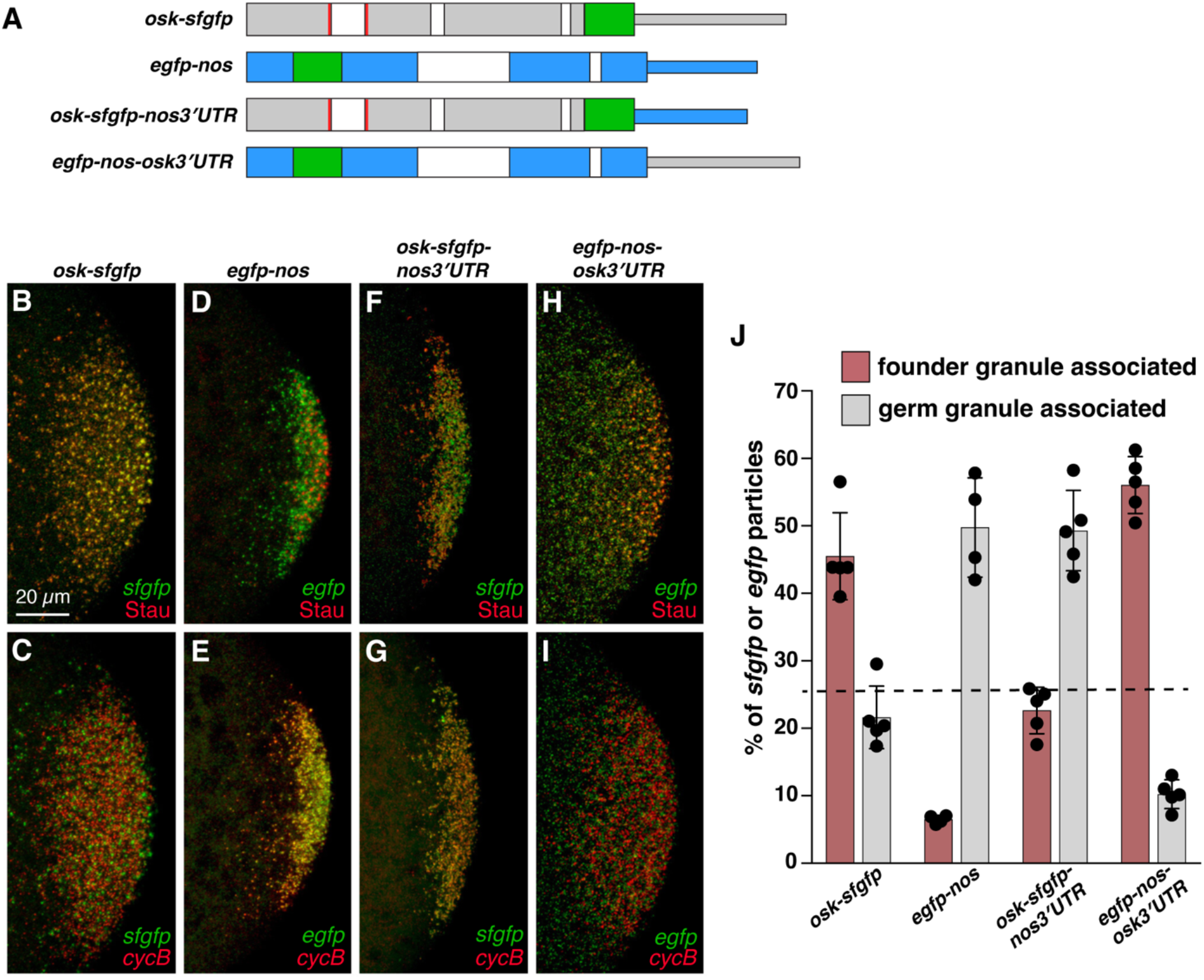
The *osk* 3’ UTR is sufficient for founder granule targeting. (A) Structure of transgenes. Gray boxes: *osk* exonic sequences; blue boxes: *nos* exonic sequences; white boxes: introns; red lines: sequences that create the SOLE; thinner boxes indicate 3’ UTRs. (B-I) Confocal z series projections of early embryos (≤ nuclear cycle 4). Transgenic RNAs are detected by FISH with probes for *sfgp* or *egfp* (green). *cycB* FISH (red) marks germ granules (B-E) and anti-Stau immunofluorescence (red) marks founder granules (F-I). Note that *cycB* is found in about 50% of all germ granules (Little et al., 2015). (J) Quantification of colocalization between transgenic mRNAs and either germ granules or founder granules; n = 4-5 embryos each. Values are mean ± standard deviation; p value for each pair <0.0001 as determined by 1-way ANOVA with Tukey’s post-hoc test and by Student’s t-test.

Based on the previous finding that fewer than 25% of *osk* particles in the germ plasm colocalize with germ granules (Little et al., 2015), we set a threshold for granule association at 25% of *sfgfp* or *egfp* particles colocalized with germ granule or founder granule markers as described above. The control *osk-sfgfp* and *egfp-nos* mRNAs behaved as expected. 43% of *osk-sfgfp* particles colocalized with founder granules (Figure 4B, J), whereas colocalization with germ granules was 22%, below the threshold (Figure 4C, J). Conversely, 50% of *egfp-nos* particles colocalized with *cycB*-containing germ granules (Figure 4E, J), similarly to the behavior of *nos* (Little et al., 2015). In contrast, colocalization of *egfp-nos* with founder granules was 6% (Figure 4D, J). Similarly to *egfp-nos*, 49% of *osk-sfgfp-nos3’UTR* particles were colocalized with germ granules (Figure 4G, J), whereas 23% colocalized with founder granules (Figure 4F, J). Furthermore, similarly to *osk-sfgfp*, 56% of *egfp-nos-osk3*’*UTR* particles colocalized with founder granules (Figure 4H, J), whereas only 10% colocalized with germ granules (Figure 4I, J). Thus, the granule preference of the transgenic mRNAs was determined by the 3’ UTRs. Taken together, these results indicate that the *osk* 3’ UTR targets mRNAs to founder granules just as the *nos* 3’ UTR targets mRNAs to germ granules.

### The *osk* coding sequences and/or 5’UTR, but not the SOLE, are required to maintain *osk* RNA accumulation in late-stage oocytes

Although the *osk* 3’UTR was sufficient to target *nos* to founder granules, *egfp-nos-osk3*’*UTR* mRNA was only weakly enriched in the germ plasm as compared to *osk-sfgfp* RNA (Figure 5A, B). To determine whether this resulted from a defect in the initial localization or a defect in maintenance of *egfp-nos-osk3*’*UTR* localization, we monitored localization over the course of late oogenesis, from stage 10-13. In contrast to *osk-sfgfp* RNA, *egfp-nos-osk3*’*UT*R did not continue to accumulate at the posterior after stage 10, consistent with the lack of the SOLE. Moreover, the amount of *egfp-nos-osk3*’*UTR* localized at the posterior of the oocyte decreased from stage 12 to stage 13, whereas *osk-sfgfp* RNA was largely unchanged (Figure 5C-H, L). Total transgenic RNA levels remained constant between these stages, suggesting that the loss of *egfp-nos-osk3*’*UTR* is unlikely a result of degradation (Figure S2). Thus, this trend is most consistent with a failure to maintain localized *egfp-nos-osk3*’*UTR* mRNA in late-stage oocytes. This result also suggests that the *osk* coding sequences or 5’ UTR contain an element important for maintenance of *osk* mRNA for the duration of oogenesis. Since the *SOLE*^*UA-mut*^ mRNA was less enriched in the germ plasm of late oocytes than *osk-sfgfp* mRNA, we hypothesized that the SOLE could be such a maintenance element. In contrast to *egfp-nos-osk3’UTR* mRNA, however, *oskSOLE*^*UA-mut*^*-sfgfp* mRNA was not lost from the posterior after stage 12 (Figure 5I-L). Thus, the SOLE most likely functions in the late-stage growth of founder granules rather than in maintaining *osk* mRNA after it is localized.

**Figure 5.**
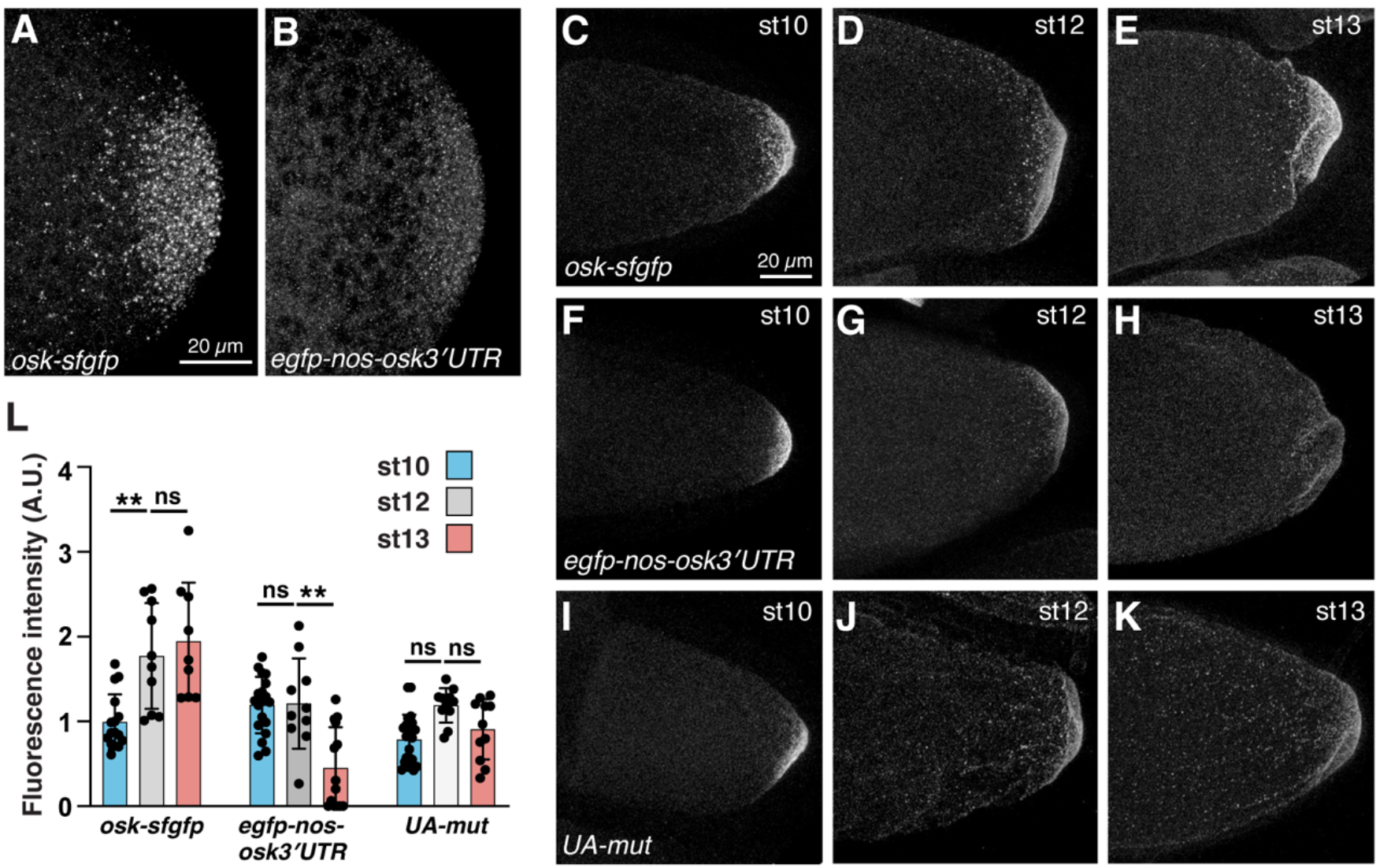
Additional features are required to maintain *osk* after stage 12. (A-B) Confocal z series projections of early embryos showing the entirety of the germ plasm (nuclear cycle 3-5). (C-K) Confocal z projections of oocytes at stage 10 (C, F, I), stage 12 (D, G, J), and stage 13 (E, H, K) capturing the entirety of the germ plasm. Transgenic RNAs were detected by FISH with probes for *sfgfp* or *egfp*. (L) Quantification of fluorescence intensity of the germ plasm from stages 10 to 13; n = 9-22 oocytes each. Values are mean ± standard deviation. ** p < 0.01 as determined by Kruskal-Wallis one-way analysis of variance, with Dunn’s post-hoc test.

## DISCUSSION

We assessed the function of the earlier acting EJC/SOLE complex in late-phase *osk* localization and found that the SOLE, but not the EJC, is required. Whereas the function of the SOLE in the earlier localization phase relies only on the structure of the proximal stem (Ghosh et al., 2012), both the sequence of the proximal stem and its structure are important for late-phase localization. How the SOLE collaborates with the EJC to promote kinesin-dependent *osk* motility and what, if any regulatory factor interacts with it are not yet known. The sequence-dependence of the SOLE and lack of requirement for the EJC in late-phase localization suggests a different mode of action, possibly through the binding of a different protein to the proximal stem and recruitment of new RNP components. The change in ovarian physiology with the onset of nurse cell dumping could lead to an exchange of proteins associated with *osk*, to inhibit kinesin-dependent motility and promote posterior entrapment.

The failure of *osk-K10_3*’*UTR* mRNA to localize at late-stages of oogenesis despite the presence of the SOLE indicates that similarly to the earlier phase, late-phase localization depends on both the SOLE and the 3’ UTR. This dependence of the 3’ UTR is independent of hitchhiking and we presume that it represents a similar requirement for 3’ UTR-binding proteins in assembling a localization-competent RNP. Both Hrp48 and Stau bind to the *osk* 3’ UTR, but only Stau remains associated with the *osk* RNP through the end of oogenesis making it a good candidate (Huynh et al., 2004; Mohr et al., 2021; Yano et al., 2004). However, the requirement for Stau in the initial posterior localization and translation of *osk* and the requirement for Osk protein to maintain *osk* mRNA at the posterior of the oocyte makes it difficult to test this prediction.

Although the germ granule mRNA *nos* and the founder granule mRNA *osk* both localize to the germ plasm during late oogenesis by diffusion and entrapment, they are assembled into distinct RNP granules in the germ plasm (Little et al., 2015; Sinsimer et al., 2011). By swapping the *osk* and *nos* 3’ UTRs, we showed that the *osk* 3’UTR contains information that directs it specifically to founder granules independently of the SOLE. Moreover, the SOLE cannot override the targeting function of the *nos* 3’UTR, suggesting that although it is required for *osk* to continue to accumulate during late-stages of oogenesis, it does not confer granule specificity. Results from swapping the *osk* and *nos* 3’ UTRs also suggest that either *osk* 5’ UTR and/or coding sequences other than the SOLE contribute to maintenance of *osk* mRNA in founder granules or to the maintenance of the granules themselves. Since multivalent interactions are typically required for inclusion of components in phase separated condensates (Banani et al., 2017), it is not surprising that binding of founder granule components to multiple sites within *osk* would be required for *osk* to be retained within granules. Further dissection of the sequence requirements and identification of interacting factors will be necessary to define the mechanisms by which the various *osk* elements accomplish the different tasks.

The process by which *osk* achieves its posterior localization is remarkably complex and labor intensive, involving distinct machineries for transport into the oocyte, movement to the posterior pole during stages 8-10, and further accumulation during late-stages of oogenesis. Given the dependence of embryonic abdominal patterning and germ cell formation on the amount of *osk* mRNA localized during oogenesis (Ephrussi and Lehmann, 1992; Gavis and Lehmann, 1994; Smith et al., 1992), the reliance on numerous distinct contributions to *osk* localization likely provides robustness to processes governing the targeting of *osk* RNPs to the right location and the accumulation of sufficient *osk* there.

## MATERIALS AND METHODS

### Construction of transgenes and transgenic lines

The *osk-sfgfp* and *osk-Δi-sfgfp* transgenes and transgenic lines were previously described and contain *sfgfp* sequences inserted just before the *osk* stop codon in an 8 kb genomic *osk* rescue fragment in the pattB vector (Eichler et al., 2020). *osk-Δi-sfgfp* lacks the first and second *osk* introns (Eichler et al., 2020). The *egfp-nos* transgenes and transgenic lines were previously described and contain *egfp* sequences inserted just after the *nos* start codon in a 4.3 kb *nos* genomic rescue fragment (Forrest et al., 2004). To generate *osk-sfgfp-nos3’UTR*, the *osk* 3’ UTR and 3’ genomic DNA was removed from the plasmid pattB-osk-sfgfp (Eichler et al., 2020) and replaced with a 1.3 kb fragment from containing the *nos* 3’ UTR and 451 bp of 3’ genomic *nos* DNA. To generate *egfp-nos-osk3’UTR*, a fragment containing the *nos* promoter and 5’ UTR, *egfp*, and *nos* coding region (including introns) was removed from pCaSpeR-Pnos-gfp-nos (Forrest et al., 2004), fused to the *osk* 3’ UTR and 3.2 kb of *osk* 3’ genomic sequences, and cloned into pattB. To generate the *osk-sfgfp* SOLE mutant transgenes, a 774 bp SphI-SacI fragment from pattB-osk-sfgfp (Eichler et al., 2020) was replaced with a 774 bp SphI-SacI fragment synthesized by Genewiz with either the PS-Lz, UA^mut^, or UA^mut^-comp mutation shown in Figure 2A. *UASp-osk-sfgfp-K10_3*’*UTR* was generated by inserting fragment from pattB-osk-sfgfp (Eichler et al., 2020) spanning a naturally occurring BamHI site just before the *osk* transcription start site to an engineered BamHI site immediately following the *osk* stop codon into the BamHI site of pattB-UASp. Transgenes were integrated into the attP40 site by phiC31-mediated recombination.

### Tissue collection and fixation

#### Ovaries

Females were fed on yeast paste at 25°C for 3 days. Ovaries were dissected into PBS, lightly teased apart, and fixed and stepped into methanol as previously described (Abbaszadeh and Gavis, 2016). Ovaries were stored in methanol at -20 °C for no more than 1 month. Ovaries used for RNA extraction were transferred to a 1.5 mL eppendorf tubes after dissection, flash frozen in liquid nitrogen, and stored at -80 °C.

#### Embryos

Embryos were collected on apple juice agar plates at room temperature, then dechorionated, fixed, and devitellinized as described (Abbaszadeh and Gavis, 2016). Embryos were stored in methanol at -20 °C for up to 1 month. Embryos used for RT-qPCR were dechorionated, flash frozen in liquid nitrogen, and stored at -80 °C.

### Single molecule fluorescence in situ hybridization (smFISH)

smFISH was performed according to Abbaszadeh and Gavis (2016). smFISH probes consisting of 20 nt oligonucleotides with 2 nt spacing were designed with Stellaris Probe Designer and synthesized by Biosearch Technologies. Probe sets complementary to *cycB* (CG3510; 48 oligos) or *egfp* (32 oligos) were conjugated to Atto 647N dye (Sigma-Aldrich) or Atto 565 dye (Sigma-Aldrich) and purified by HPLC as previously described (Raj et al., 2008). smFISH probes of 20 nt oligonucleotides with 2 nt spacing for *sfgfp* (31 oligos) were designed with Stellaris Probe Designer and purchased already conjugated to Quasar 670 fluorophore from Biosearch Technologies. For quantitation of total localized fluorescence intensity, 1 µL of probes per 100 µL of hybridization buffer was used; for colocalization analysis 3 µL of probes per 100 µL was used. Ovaries or embryos samples were mounted under #1.5 glass coverslips (VWR) in Vectashield Mountant (Vector Laboratories) for quantification of total localized fluorescence intensity.

### Immunofluorescence

Immunofluorescence was performed as describe in Eichler et al. (2020). Embryos were incubated in rabbit ant-Staufen #36.2 (kindly provided by D. St Johnston) diluted 1:2000 in PBHT (PBS, 0.1% Tween-20, 0.25 mg/ml heparin [Sigma-Aldrich], 50 mg/ml tRNA [Sigma-Aldrich]) overnight at 4 °C with rocking. Alexa-488 goat anti-rabbit secondary antibody (Molecular Probes) was diluted 1:1000 in PBHT and applied for 2 hr at room temperature with rocking. Embryos were mounted as described above. For double immunofluorescence/smFISH experiments, immunostaining was performed first, then embryos were refixed in 4% PFA for 30 min at room temperature with rocking and rinsed 4× with PBST before proceeding with smFISH as described above. Embryos were mounted in Prolong Diamond Antifade Mountant (Thermo Fisher Scientific).

### Fluorescence intensity quantification

Confocal imaging for all experiments except for Figure 3 was performed using a Leica SP5 laser scanning microscope with a 63× 1.4 NA oil immersion objective and GaAsP “HyD” detectors in standard detection mode. For the smFISH experiment in Figure 3, confocal imaging was performed using a Nikon A1 microscope with a 40x 1.3NA oil immersion objective and GaAsP detectors. All imaging parameters were kept identical within each experiment. For quantification of total fluorescence intensity, z-series with a 2 µm step size were used to capture the entire germ plasm-localized signal. Image processing and analysis were done in FIJI. Z-projections were made with the “sum slices” function and the threshold adjusted so the entire localized signal was included. The total fluorescence intensity of the localized signal (integrated density function in FIJI) was then measured. For the time series analysis, an ROI was drawn to encompass the germ plasm and fluorescence intensity was measured for the ROI, then the ROI was moved to an anterior region of the oocyte and background fluorescence intensity was measured and subtracted from the germ plasm ROI measurement. Data are displayed as mean ± standard deviation. For 2-sample comparisons, the Student’s t-test was used; for multiple comparisons, a one-way ANOVA was used with either Tukey or Dunnett’s post-hoc test as indicated in the figure legend. Statistical analysis was performed using GraphPad Prism software.

### RT-qPCR

RNA was extracted from dechorionated embryos using the RNeasy kit (Qiagen). 0.75 µg total mRNA was used to generate cDNA using the Quantitect RT kit (Qiagen). 2 µl cDNA was combined with 25 µl 2× TaqMan Gene Expression Master Mix (Thermo Fisher Scientific), 2.5 µl of 20× TaqMan Gene Expression Assay (Thermo Fisher Scientific, *sfgfp* custom – APT2CRZ, 4331348, *egfp* Mr 04097229_mr Enhance, or *rpl7* Dm 01817653, 4351372), and 20.5 µl of nuclease free H_2_O. qPCR was performed on an Applied Biosystems 7900HT standard 96-well qPCR instrument. Three biological replicates were performed with three technical replicates each, all using a CT threshold of 0.6613619. Technical replicates were averaged and the three biological replicates were normalized to the *rpl7* control using the ΔC_t_ method and presented as mean ± standard deviation. Statistical significance was determined by Student’s t-test or by one-way ANOVA and Tukey’s multiple comparisons tests, as indicated in the Figure legends, using GraphPad Prism software.

## ACKNOWLEDGEMENTS

We are grateful to D. St Johnston for anti-Stu antibody and A. Ephrussi and P. Macdonald for plasmid DNA. We thank G. Laevsky and S. Wang for assistance with confocal microscopy, S. Chatterjee for technical assistance, and A. Hakes for comments on the manuscript. This work was supported by National Institute of Health (NIH) grant R35 GM126967 to ERG. CEE was supported by NIH training grant T32 GM007388.

---

**Figure S1.**
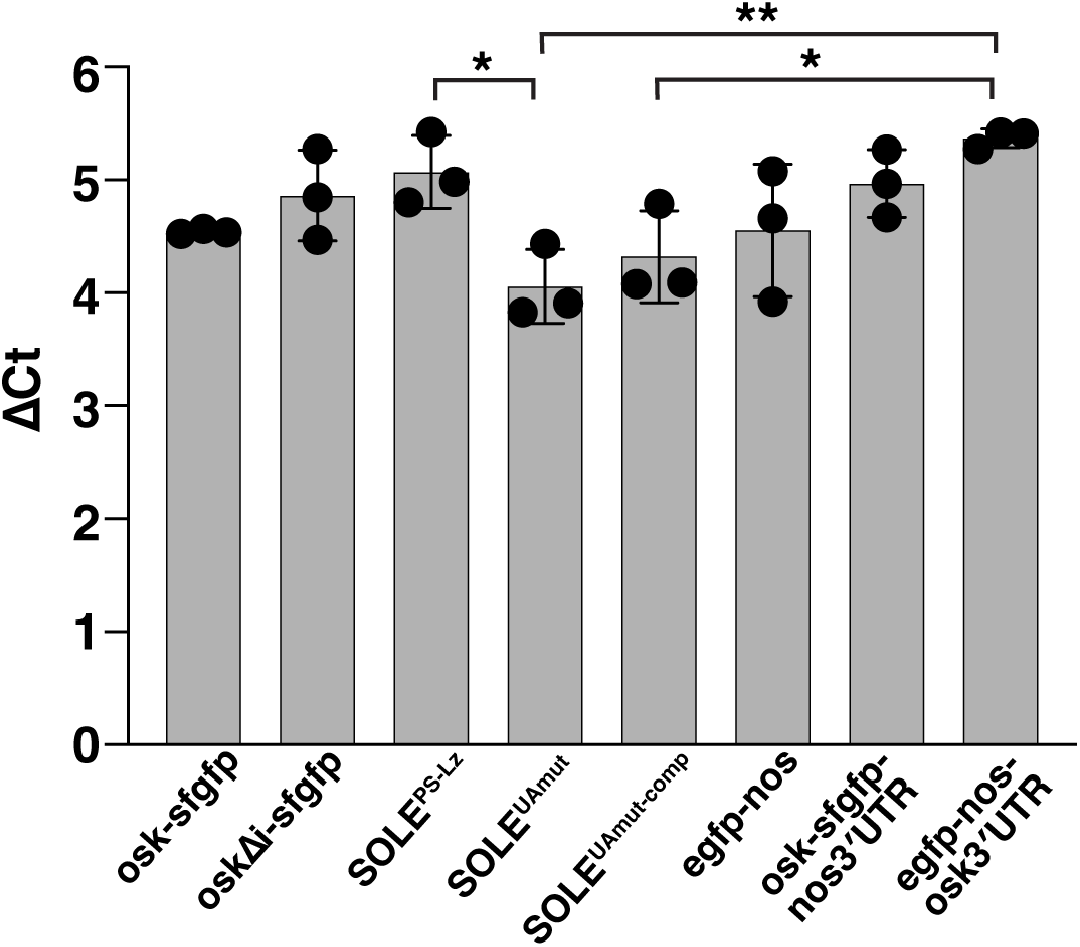
Quantitation of transgenic RNA levels. RT-qPCR quantification of transgenic RNA extracted from early embryos, normalized to *rpl7* RNA. * p < 0.05; p** p < 0.01, as determined by one-way ANOVA and Tukey’s multiple comparisons test.

**Figure S2.**
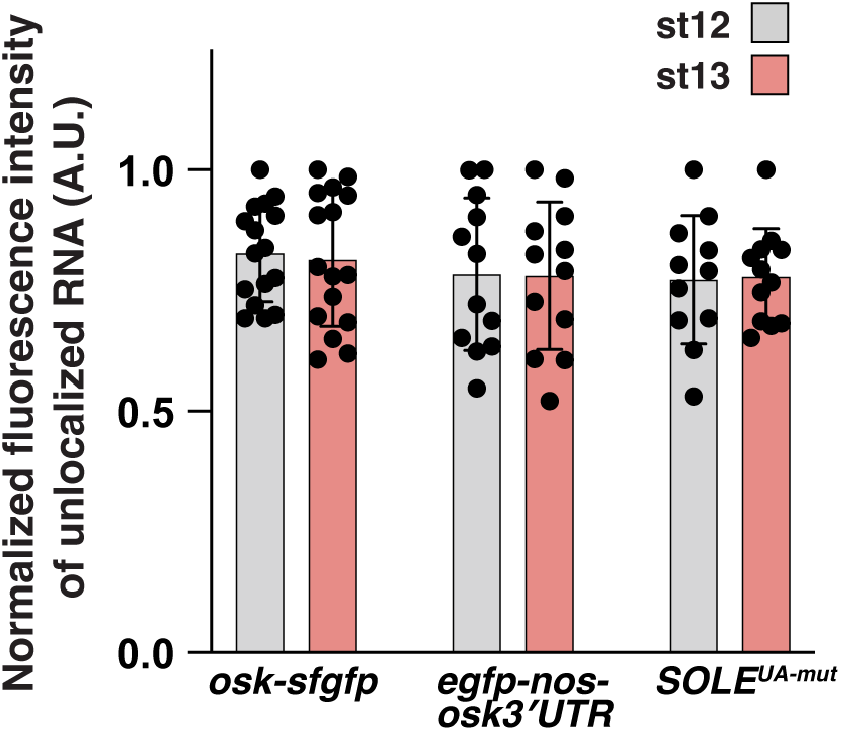
Stability of transgenic RNAs during late-stages of oogenesis. Fluorescence intensity measurements of unlocalized *osk-sfgfp, egfp-nos-osk3’UTR* and *osk-sfgfpSOLE*^*UA-mut*^ mRNAs in stage 12 and stage 13 oocytes. n = 11-14 oocytes each. The values are not statistically significant as determined by one-way ANOVA and Tukey’s post-hoc test.

